# Accurate and scalable multi-disease classification from adaptive immune repertoires

**DOI:** 10.1101/2025.08.12.669991

**Authors:** Natnicha Jiravejchakul, Ayan Sengupta, Songling Li, Debottam Upadhyaya, Mara A. Llamas-Covarrubias, Florian Hauer, Soichiro Haruna, Daron M. Standley

## Abstract

**Background:** Machine learning models trained on paratope-similarity networks have shown superior accuracy compared with clonotype-based models in binary disease classification. However, the computational demands of paratope networks hinder their use on large datasets and multi-disease classification.

**Methods:** We reanalyzed publicly available T cell receptor (TCR) repertoire data from 1,421 donors across 15 disease groups and a large control group, encompassing approximately 81 million TCR sequences. To address computational bottlenecks, we replaced the paratope-similarity network approach (Paratope Cluster Occupancy or PCO) with a new Fast Approximate Clustering Techniques (FACTS) pipeline, which is comprised of four main steps: (1) high-dimensional vector encoding of sequences; (2) efficient clustering of the resulting vectors; (3) donor-level feature construction from cluster distributions; and (4) gradient-boosted decision tree classification for multi-class disease prediction.

**Findings:** FACTS processed 10^7^ sequences in under 120 CPU hours. Using only TCR data, and evaluated with 5-fold cross-validation, it achieved a mean ROC AUC of 0.99 across 16 disease classes. Compared with the recently reported Mal-ID model, FACTS achieved higher donor-level classification accuracy for BCR (0.840 vs. 0.740), TCR (0.882 vs. 0.751), and combined BCR+TCR datasets (0.904 vs. 0.853) on the six-class Mal-ID benchmark. FACTS also preserved biologically meaningful signals, as shown by unsupervised t-SNE projections revealing distinct disease-associated and potentially age-associated clusters.

**Interpretation:** Paratope-based encoding with FACTS-derived features provides a scalable and biologically grounded approach for adaptive immune receptor (AIR) repertoire classification. The resulting classifier achieves superior multi-disease diagnostic performance while maintaining interpretability, supporting its potential for clinical and population-scale health profiling.

**Funding:** This study was supported by the Japan Society for the Promotion of Science (JSPS) KAKENHI [JA23H034980], the Japan Agency for Medical Research and Development (AMED) [JP25am0101001], and the Kishimoto Foundation Fellowship.

**Research in context:** *Evidence before this study:* T and B cell receptor (TCR and BCR) repertoires encode lifelong immunological memory and antigen-specific responses, making them valuable biomarkers for disease diagnosis and prediction. Existing machine learning (ML) models for adaptive immune receptor (AIR) repertoires often rely on clonotype-based representations, which limit shared receptor detection between donors and thus reduce cross-individual disease signature detection. Most models also lack robust multi-disease, population-scale performance. Our previous work showed that representing repertoires as paratope-similarity networks increased the fraction of shared receptors between donors and improved disease classification. However, their computational complexity has limited their scalability for the large datasets required in multi-disease classification.

*Added value of this study:* We introduce FACTS, a unified ML framework integrating paratope similarity with scalable sequence encoding. Applied to TCR repertoires from 1,421 donors across 15 diseases and one control group, FACTS maintained high performance while efficiently processing 81 million sequences on standard CPU infrastructure. Compared to Mal-ID, our paratope-encoded method achieved significantly higher donor-level accuracy and revealed biologically meaningful disease- and potentially age-associated patterns.

*Implications of all the available evidence:* FACTS offers high accuracy, and interpretability for multi-disease classification, bringing AIR repertoire-based diagnostics closer to clinical translation and potentially guiding precision immunotherapy and immune-based therapeutic discovery for a wide range of diseases.

## 1 Introduction

B and T lymphocytes generate diverse antigen receptors through somatic V(D)J gene recombination^1,2^. The resulting B-cell receptors (BCRs) and T-cell receptors (TCRs), collectively known as adaptive immune receptors (AIRs), record a lifelong molecular record of antigen encounters. AIRs can be profiled through a routine blood draw, followed by standard sequencing, offering a minimally invasive, hypothesis-free source of disease biomarkers that can detect immune perturbations before clinical symptoms appear. AIR repertoire analysis, therefore, holds promise for early and personalized disease detection, prognosis, and the discovery of immune-based therapeutics^3,4^.

In recent years, machine learning (ML) has emerged as a powerful approach for identifying disease-specific signatures from AIR repertoires, as reviewed by Mhanna *et al*.^5^. While binary classifiers have been effective in distinguishing cohorts of patients with specific disease from healthy donors, such models do not reflect the complexity of the immune system, which responds to diverse pathogens and disease conditions. Multi-disease classification, as exemplified by the recently introduced Mal-ID program on 6 disease/healthy classes, is thus both more realistic and clinically relevant than previous models^6^. However, training such ML models requires larger and more heterogeneous datasets, increasing computational demands and making efficiency a critical factor for clinical translation.

Most existing ML models rely on clonotype-based repertoire representations in which sequences that share the same V-gene and possess identical, or highly similar, complementarity-determining region 3 (CDR3) sequences are grouped together^3,7^. Due to the immense diversity of BCRs and TCRs, the proportion of clonotypes shared between donors is extremely low. Studies report approximately 1-6% overlap in IGH clonotypes and around 8-13% in TCRβ clonotypes across healthy donors^8,9^. This sparsity limits the generalizability and sensitivity of clonotype-centric ML approaches.

Paratope-based representations address this limitation by focusing on receptor residues likely to interact with antigens or peptide-MHC complexes^10,11^. This avoids combining categorial germline gene annotations with continuous amino acid sequences and extends analysis beyond CDR3 to include CDR1 and CDR2, enabling statistically robust sequence comparisons. Notably, paratope clusters are typically antigen- and epitope-specific. In prior work, grouping BCRs by paratope similarity (80% sequence identity and 90% alignment coverage) yielded clusters in which more than 95% of antibodies recognized the same antigen, with epitope-specific neutralizing antibodies co-clustering as well^12^. Paratope similarity networks were shown to provide superior features for binary disease classification, outperforming state-of-the-art clonotype-based models such as DeepRC and immuneML^13^. However, constructing such networks requires all-against-all similarity comparison, limiting scalability to datasets with fewer than 10^6^ sequences.

In this study, we developed a scalable paratope-based feature construction pipeline to enable multi-disease classification across large AIR datasets. We replaced computationally intensive paratope similarity graphs with an efficient sequence encoding and clustering strategy called Fast Approximate Clustering Techniques (FACTS). To validate FACTS, we reanalyzed publicly available TCR repertoire data from 1,421 donors across 15 disease groups and one large control group. Compared to our previous method, Paratope Cluster Occupancy (PCO), FACTS dramatically reduced computational time while maintaining high classification performance, achieving a mean five-fold ROC AUC of 0.99. On the six-class Mal-ID benchmark, FACTS using only TCR data achieved an accuracy of 88.2%, which is slightly higher than the 85.3% accuracy of the Mal-ID model that used both BCR and TCR data. Moreover, unsupervised embedding projections revealed disease- and demographic-associated clusters, supporting FACTS as a robust, scalable, and interpretable framework for repertoire-based diagnostics with potential for clinical and population-scale applications.

## 2 Methods

### Dataset curation and preprocessing

Adaptive immune receptor (AIR) sequencing data from a total of 1,421 donors were obtained from publicly available studies spanning 16 disease groups, including healthy individuals, patients with autoimmune disease, infectious disease, cancer, and vaccinated donors. Data for healthy donors, human immunodeficiency virus (HIV) infection, influenza vaccination, coronavirus disease 2019 (COVID-19), systemic lupus erythematosus (SLE), and type 1 diabetes (T1D) were sourced from the Mal-ID public dataset^6^ (Supplementary Table S1). Samples labeled as “HIV-negative” were excluded from the healthy donor group in our study. Data for the remaining 10 disease groups, as well as additional healthy donors, were obtained from 12 published studies^14-25^ (Supplementary Table S1). No original sequencing data were generated in this study, and ethical approval for secondary analysis was not required by the authors’ institution.

### Paratope extraction

In the first step of the FACTS pipeline, paratope regions were extracted from raw AIR sequences using ANARCI^26^, with residue numbering assigned according to the IMGT scheme^27^. For each BCR and TCR sequence, paratopes were defined as all positions within CDR1, CDR2, and CDR3, concatenated to form a single pseudo-sequence used in all downstream encoding and clustering steps.

### Sequence encoders

Following paratope extraction, each pseudo-sequence was transformed into a fixed length embedding using a protein language model. An encoder for TCR sequence was developed by adopting a Transformer-based encoder-only architecture trained on large-scale, publicly available AIR datasets. Training employed the masked language modelling (MLM) objective from BERT^28^, in which random amino acid tokens are masked, and the model is optimized to predict the missing residues. For the TCR encoder model, weights were initialized from ESM-2^29^ and fine-tuned on TCR datasets using paratope-focused sequence inputs. The resulting “MyImmune” TCR encoder generated dense vector embeddings for each paratope pseudo-sequence, which were subsequently used for clustering and classification. By default, FACTS uses IgBert^30^ for encoding BCR sequences.

### Clustering of sequence embeddings

To group similar paratope embeddings, MiniBatchKMeans, a memory- and time-efficient variant of k-means, was applied using the scikit-learn^31^ implementation with k ranging from 100 to 1,000. A batch size of 1,024 was used, and initial centroids were seeded with the k-means++ initializer to improve stability. Clustering quality was assessed across the k range using inertia (within-cluster sum of squares) and silhouette coefficient scores^32^. MiniBatchKMeans scales approximately as O(N log k) per pass through the dataset, enabling efficient clustering of tens of millions of paratope embeddings without prohibitive computational cost.

### Construction of donor-level features

Following clustering, donor-level feature vectors were generated by aggregating cluster membership data. For each donor, the frequency distribution of clustered paratope embeddings was computed as the proportion of sequences assigned to each cluster. Inter-cluster mutual information was also calculated. This produced a fixed-length vector representing each donor’s immune repertoire, which served as input for downstream classification. Mathematical details of how the donor-level features are defined, was reported in the previous study^13^.

### Classification models and statistical analysis

All classification models were trained using the XGBoost^33^ implementation in soft probability mode. Model performance was assessed using five-fold cross-validation with donor-level stratification to prevent donor overlap between folds. Evaluation metrics included accuracy, ROC AUC, macro-averaged F1 score (macro F1), sensitivity, and specificity. Paired AUC values were compared using the Wilcoxon signed-rank test. Ninety-five percent confidence intervals (CIs) for performance metrics were calculated via bootstrap resampling (n = 10,000 iterations).

### Benchmarking sequence encoders

MyImmune TCR encoder was benchmarked against several open-source protein and immune receptor language models, including TCRBert^34^, ESM-2^29^, ProtTrans^35^, SCEPTR^36^, and UniTCR^37^. For BCR sequences, six publicly available encoders; IgBERT^30^, AntiBERTa2^38^, AntiBERTy^39^, ESM-2^29^, ProtTrans^35^, and AbLang^40^, were evaluated. Details of each encoder are provided in Supplementary Table S3. Each model was integrated into our pipeline using identical clustering and classification procedures, differing only in the embedding step. Benchmarking was performed on TCR and BCR datasets from the Mal-ID cohort^6^ (499 donors for TCR, and 550 donors for BCR), across five disease classes and one healthy control group.

## 3 Results

### Scalable paratope-based classification pipeline

We developed a scalable machine learning framework to classify AIR repertoire data using paratope-derived sequence features. Previously, these features were computed by combining paratope clusters with a paratope similarity graph, denoted Paratope Cluster Occupancy (PCO)^13^ (Figure 1a). To avoid the all-against-all comparisons required for graph construction, we modified the workflow: sequences were first encoded as vectors, then efficiently clustered using these vector representations. We term this approach Fast Approximate Clustering Techniques (FACTS) (Figure 1b).

**Figure 1.**
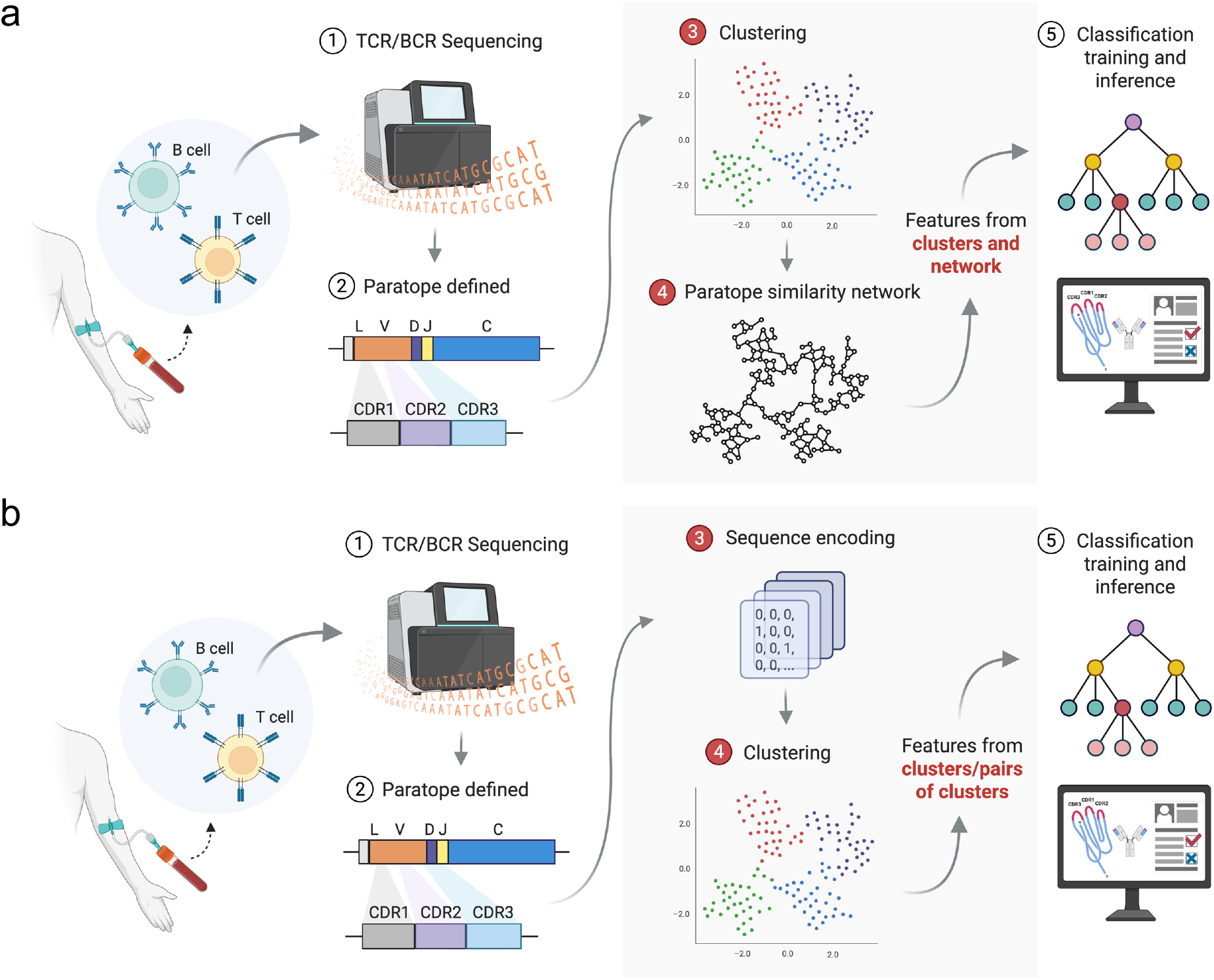
Scalable paratope-based classification pipelines for immune repertoire data. (**a**) The original paratope cluster occupancy (PCO) framework. After TCR/BCR sequencing, paratope regions (CDR1, CDR2, and CDR3) are defined and clustered. A paratope similarity network is constructed via all-against-all comparisons of paratope sequences, and donor-level features are derived from both clusters and networks. These features are then used to train a classification model. (**b**) The proposed fast approximate clustering techniques (FACTS). Instead of pairwise comparison, paratope sequences are first encoded into vector embeddings before clustering. Donor-level features are constructed from cluster frequencies and inter-cluster relationships. This reordering avoids the computational cost of paratope graph construction and enables scalable training of classification models.

The pipeline begins with the extraction of paratope regions from BCR or TCR sequences, represented as pseudo-sequences composed of the concatenated CDR1, CDR2, and CDR3. These sequences are encoded, and the resulting embeddings are clustered using k-means (k = 50-500). A silhouette coefficient score^32^ is used to determine a good starting point for choosing the number of clusters k. Donor-level feature vectors are constructed from cluster frequency distributions and inter-cluster mutual information. These features are then used to train gradient-boosted decision tree models for multi-class disease classification.

The use of FACTS qualitatively improved the scalability compared with PCO (Figure 2). Two PCO implementations were tested: one using an exact all-against-all comparison and one using an MMSEQS^41^-based approximation^13^. Even with MMSEQS acceleration, PCO runtimes increased quadratically with input size (R^2^ = 0.9997). In contrast, FACTS scaled linearly when using a small number of clusters: at k = 50, runtime increased to 115 ms per 1,000 sequences (R^2^ = 0.9957); at k = 500, runtime rose to 1,158 ms per 1,000 sequences (R^2^ = 0.9195), reflecting the added cost of inter-cluster comparisons. Processing 5 million sequences required approximately 1.9 CPU-hours using FACTS, compared to about 324 CPU-hours with MMSEQS-accelerated PCO. This demonstrated an approximate 170-fold improvement in efficiency and scaled linearly with increasing data size.

**Figure 2.**
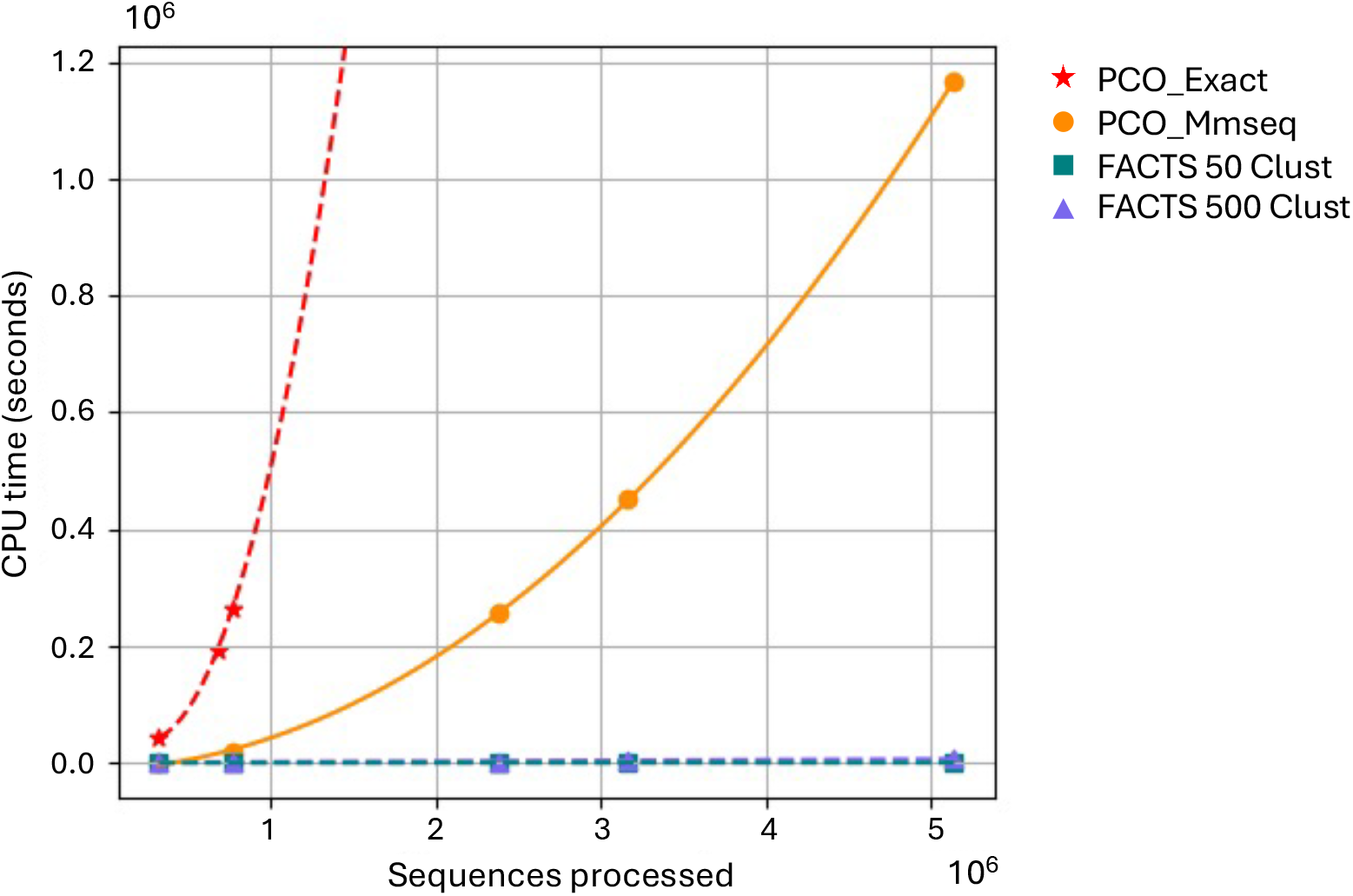
Runtime comparison between PCO and FACTS pipelines. CPU time (in seconds) is plotted against the number of sequences processed for four methods. The original PCO pipeline using exact paratope similarity (PCO_Exact, red stars) shows quadratic runtime scaling with input size due to its all-against-all comparison step. A faster variant using MMseqs2 (PCO_MMSeqs, orange circles) improves efficiency but still scales nonlinearly. In contrast, the FACTS approach achieves near-linear scaling, with FACTS configured for 50 clusters (blue squares) and 500 clusters (purple triangles) showing minimal runtime increase over 5 million sequences. The FACTS pipeline achieves up to ∼170-fold speedup compared to the exact PCO method, making it suitable for large-scale repertoire analysis.

### Robust multi-disease TCR repertoire classification achieved with encoded paratope features

To assess classification performance, we reanalyzed publicly available TCR repertoire data comprising 1,421 donors across 16 disease classes (Supplementary Table S1). The classifier achieved a mean ROC AUC of 0.9885 in five-fold cross-validation (Figure 3a; Supplementary Table S2). Per-disease AUCs were consistently high, with the lowest observed for healthy controls (0.95), while all other disease classes were greater than 0.98 (Figure 3b). The model obtained an accuracy of 81.77% (0.8177 ± 0.0090) with high specificity (0.9866 ± 0.0007) and moderately high sensitivity (0.7785 ± 0.0203) (Figure 3c; Supplementary Figure S1).

**Figure 3.**
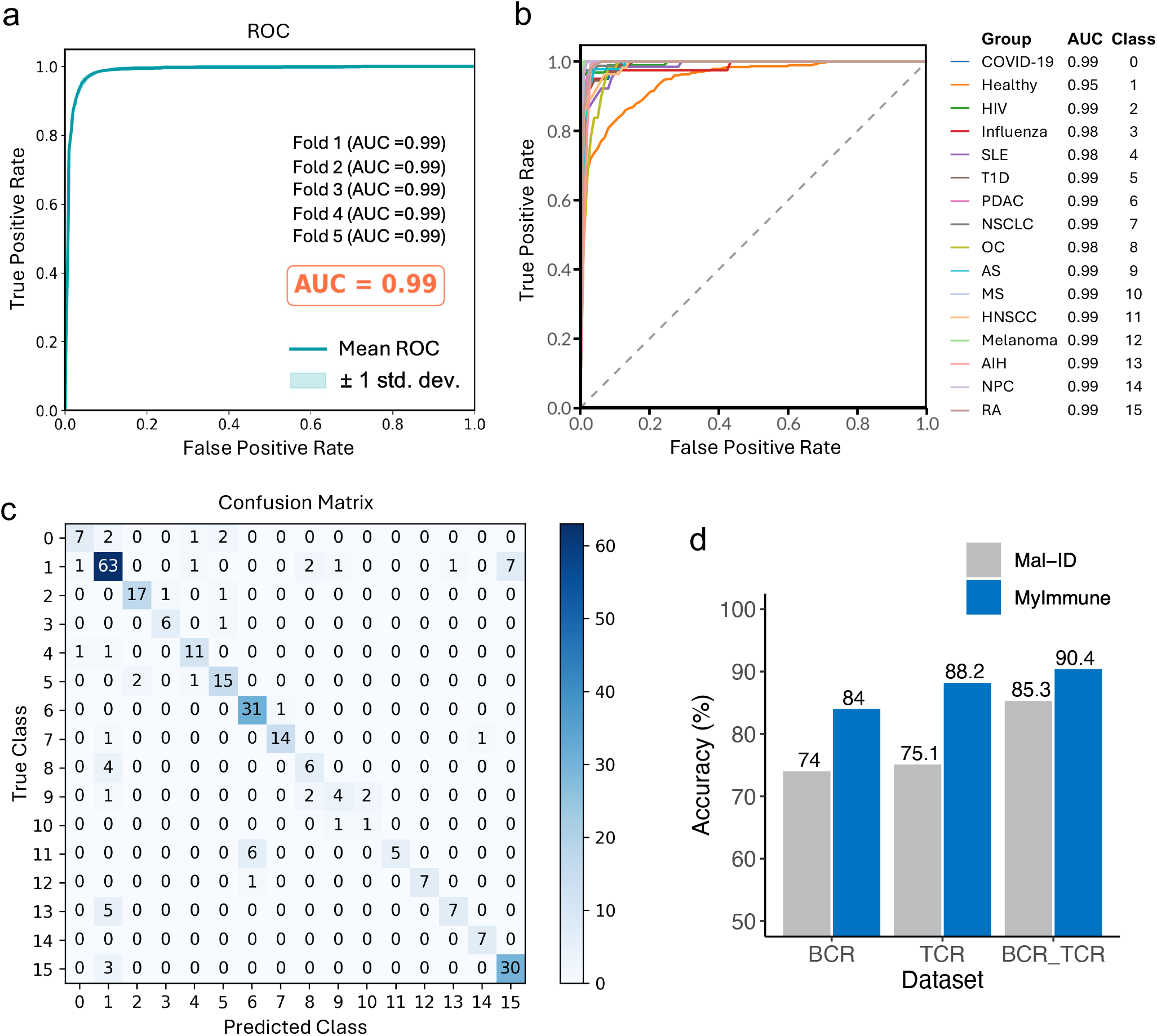
Encoded paratope features enable accurate multi-disease classification of TCR repertoires. (**a**) Receiver operating characteristic (ROC) curve showing overall performance of the paratope-based classifier on a 1,421-donor dataset comprising 16 disease classes, including healthy controls. The model achieved a mean ROC AUC of 0.99 with consistent performance across five cross-validation folds. (**b**) One-vs-rest ROC curves for each individual class showing a robust performance across all disease classes. **(c)** Confusion matrix from one of the five cross-validation folds, illustrating donor-level classification results across the 16 disease classes. Disease labels corresponding to each class ID are provided in Supplementary Table S1. **(d)** Comparative performance between our paratope-based model (MyImmune) and the original clonotype-based Mal-ID framework on the Mal-ID cohort, consisting of 550 donors for BCR and 499 donors for TCR dataset across six disease classes. Accuracy was computed as donor-level classification accuracy, representing the proportion of correctly classified donors.

We further evaluated FACTS on the Mal-ID cohort to enable a direct comparison with the Mal-ID framework (499 donors for TCR and 550 donors for BCR dataset, five disease classes, and one healthy control group). We trained separate models for each input type; BCR, TCR, and combined BCR+TCR. Using overall donor-level accuracy, our method outperformed Mal-ID for all input types: BCR (84% vs 74%), TCR (88.2% vs 75.1%), and combined BCR+TCR (90.4% vs 85.3%) (Figure 3d). ROC AUCs for the Mal-ID subset were as follows: 0.9718 (BCR), 0.9781 (TCR), and 0.9824 (BCR+TCR) (Supplementary Figure S2). Importantly, our TCR-only model achieved accuracy comparable to Mal-ID’s combined-input model. This represents a high-impact reduction in input requirements for FACTS. By relying only on TCR data, equivalent performance can be obtained with less data, lowering both the cost and complexity of diagnosis in practical use cases.

### Benchmarking dense encoders for multi-disease classification

We next evaluated the performance of our in-house trained TCR encoder, along with multiple publicly available sequence encoders, on the MAL ID dataset. For this evaluation, we varied only the encoder, keeping the rest of the classification pipeline unchanged. For BCR only models, the pipeline with IgBERT as sequence encoder achieved the highest performance (accuracy: 82.0% ± 3.7%; ROC AUC: 0.9736 ± 0.0092), outperforming Antiberta2, AntiBERTy, ESM-2, ProtTrans, and AbLang. For TCR only models, the training pipeline with our TCR encoder achieved the highest performance (accuracy: 0.8819 ± 0.0562; ROC AUC: 0.9781 ± 0.0111; F1 score: 0.8576 ± 0.0644), outperforming TCRBert, ProtTrans, ESM-2, and SCEPTR (Table 1). These results highlight the strength of dense encoders for immune receptor modeling and demonstrate that our TCR encoder establishes a strong new baseline for this task.

**Table 1.** Sequence encoders form multi-disease classification. Accuracy, AUROC, and F1 score (mean ± standard deviation) are shown for each encoder using the FACTS pipeline on the Mal-ID cohort (550 donors for BCR and 499 donors for TCR dataset across 6 disease classes).

| BCR Encoders | Accuracy | AUROC | F1 score |
| --- | --- | --- | --- |
| **IgBert** | **0.8400 ± 0.0306** | **0.9718 ± 0.0073** | **0.8386 ± 0.0386** |
| Antiberta2 | 0.7920 ± 0.0497 | 0.9693 ± 0.0080 | 0.7809 ± 0.0241 |
| AntiBERTy | 0.7500 ± 0.0173 | 0.9555 ± 0.0108 | 0.7546 ± 0.0132 |
| ESM-2 8M | 0.7440 ± 0.0522 | 0.9501 ± 0.0158 | 0.7407 ± 0.0526 |
| ProtTrans | 0.7360 ± 0.0371 | 0.9537 ± 0.0108 | 0.8057 ± 0.0514 |
| ESM-2 150M | 0.7320 ± 0.0228 | 0.9473 ± 0.0063 | 0.7281 ± 0.0369 |
| ESM-2 650M | 0.7000 ± 0.0283 | 0.9386 ± 0.0076 | 0.7031 ± 0.0366 |
| AbLang | 0.6940 ± 0.0207 | 0.9398 ± 0.0030 | 0.6880 ± 0.0403 |

| TCR Encoders | Accuracy | AUROC | F1 score |
| --- | --- | --- | --- |
| **MyImmune Encoder** | **0.8819 ± 0.0562** | **0.9781 ± 0.0111** | **0.8576 ± 0.0644** |
| TCRBert | 0.8578 ± 0.0453 | 0.9735 ± 0.0108 | 0.8201 ± 0.0533 |
| ESM-2 8M | 0.8439 ± 0.0552 | 0.9755 ± 0.0104 | 0.8153 ± 0.0586 |
| ProtTrans | 0.8438 ± 0.0458 | 0.9715 ± 0.0110 | 0.8057 ± 0.0514 |
| SCEPTR: Large model | 0.8438 ± 0.0375 | 0.9747 ± 0.0121 | 0.8135 ± 0.0425 |
| ESM-2 150M | 0.8318 ± 0.0595 | 0.9742 ± 0.0103 | 0.7969 ± 0.0673 |
| SCEPTR: Beta-chain only model | 0.8317 ± 0.0282 | 0.9727 ± 0.0105 | 0.7905 ± 0.0241 |
| SCEPTR: Default model | 0.8238 ± 0.0452 | 0.9709 ± 0.0111 | 0.7902 ± 0.0597 |
| UniTCR | 0.6875 ± 0.0594 | 0.9299 ± 0.0121 | 0.6163 ± 0.0777 |

### Encoded paratope representations capture biologically meaningful structures of TCR repertoires

To assess biological relevance, we visualized donor-level embeddings from encoded TCR paratope sequences using t-SNE. In the full dataset (1,421 donors, 16 disease classes), projections showed clear separation between disease groups, indicating that paratope-based features capture disease-specific immune signatures (Figure 4a; Supplementary Figure S3). An interactive 3D tSNE plot is provided as Supplementary Table S4. From this analysis, a batch effect was observed in the autoimmune hepatitis (AIH) dataset (Supplementary Figure S4), likely reflecting differences in experimental protocols among publicly available studies, as detailed in Supplementary Note S1.

**Figure 4.**
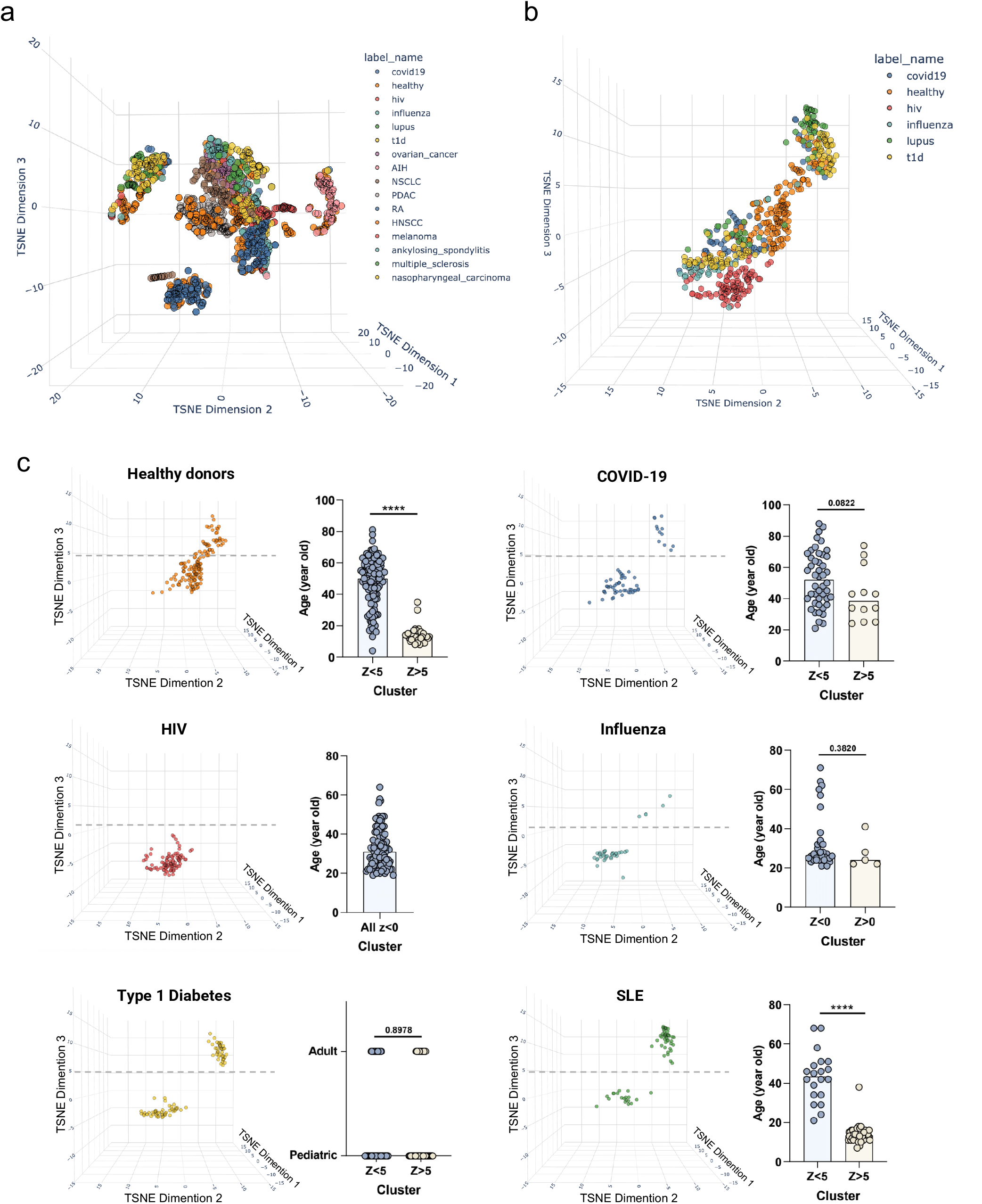
Encoded TCR paratope representations reveal disease- and age-associated structure. (**a**) t-SNE projection of donor-level TCR paratope embeddings across 1,421 donors from 16 disease classes. (**b**) t-SNE projection of the encoded TCR paratope sequences from the Mal-ID cohort at donor level (n = 499 donors, 6 disease classes). (**c**) Age association across clusters for each disease group in the Mal-ID cohort. Donors clustered along the third t-SNE axis (Z) based on age, with older individuals enriched in the Z < 5 region and younger donors in Z > 5. Statistical comparisons between clusters were performed using unpaired t-tests; **** indicates p < 0.0001.

Restricting the analysis to the Mal-ID TCR subset (n = 499 donors), which was collected and processed using a standardized protocol, obvious batch effects were reduced and while strong disease group separation was maintained (Figure 4Bb; Supplementary Table S4). Our model consistently achieved high classification accuracy and ROC AUC values when applied to either the full 1,421-donor dataset or the Mal-ID subset, demonstrating the robustness of the approach across diverse data sources (Supplementary Table S2).

Beyond disease classification, the learned feature space preserved biologically meaningful structures. Unsupervised t-SNE projection revealed a clustering pattern suggestive of age associations, particularly among SLE and healthy donors, where older individuals (Z < 5) and younger individuals (Z > 5) clustered separately. Statistically significant age differences were found within both healthy controls and SLE donors (Figure 4c). Similar, although not statistically significant, trends were observed in other disease groups, suggesting that paratope features capture clinically relevant immune signatures beyond discrete disease classification.

## 4 Discussion

In the field of immunology, ML has primarily been applied to molecular-level problems such as MHC allele prediction^42^ and TCR clustering^43^. In the latter case, paratopes, comprising the CDR1, CDR2, and CDR3 regions, represent AIR residues capable of directly interacting with antigens. Previous work has shown that paratope sequences can reflect convergent immune responses across individuals^13^, and that BCR clusters are highly antigen- and epitope-specific^12^. Other studies have demonstrated that antibodies binding the same epitope converge in paratope space and can be identified using deep learning predictors such as ParaPred^44^ and structure-aware tools like Ab-Ligity^45^.

More recently, ML methods have been applied to systems-level problems, including disease classification from AIR repertoire data. Previously, we showed that paratope networks enabled robust binary disease classification, outperforming clonotype-based methods. However, paratope networks require extensive computational resources, precluding their application to larger datasets. In the present study, we introduce FACTS, a much more scalable, framework for multi-disease classification using AIR sequences. By reversing the order of clustering and similarity comparison in our previous pipeline, we improved training runtime without apparent loss in accuracy. Rather than computing pairwise similarities first, we encoded sequences into vector representations, then performed clustering in this reduced space. We also introduced our own MyImmune TCR paratope encoder, which out-performed alternatives in the context of disease classification. The embeddings, generated using protein language models trained on natural amino acid sequences, captured biochemical and structural properties. Such embeddings have recently been used in various immune repertoire analyses^30,34,38,39,45^. MiniBatchKMeans clustering was then applied to group encoded paratope sequences, enabling efficient processing of up to hundreds of millions of sequences. Donor-level features were derived from cluster frequency distributions and inter-cluster mutual information, which were subsequently used for classification.

Benchmarking demonstrated that the MyImmune TCR paratope encoder further enhanced classification performance. Compared with multiple publicly available TCR protein language models, MyImmune consistently achieved the highest accuracy in classification of MAL-ID dataset, reinforcing the value of paratope-aware representation learning in repertoire-based diagnostics. Notably, using only TCR input, FACTS achieved accuracy comparable to a Mal-ID model that used both TCR and BCR input. This finding further suggests that reliable classification may be achievable using data from only one receptor type, which could significantly reduce diagnostic costs. Taken together, the integration of FACTS and the MyImmune encoder produced a fast, scalable, and highly accurate pipeline for immune repertoire-based disease classification.

When applied to a larger TCR dataset of 1,421 donors spanning 16 disease classes, the FACTS framework demonstrated consistently strong classification performance, although batch effects were in the AIH dataset, which likely reflects variability in experimental protocols across the cohorts. Despite this, FACTS achieved high classification accuracy not only in the large, heterogeneous dataset, but also in the smaller, more homogeneous dataset where experimental protocols were more consistent and batch effects were minimal. This consistency across diverse conditions indicates that our model is robust against batch effects. These findings emphasize the importance of validating diagnostic models using well-matched, prospectively collected clinical datasets under standardized conditions to ensure reliability in real-world applications. Overall, our findings show that paratope-encoded representations, when combined with the FACTS pipeline and MyImmune encoder, enable accurate, scalable, and biologically meaningful classification of immune repertoires, supporting their potential for clinical translation and population-scale immune profiling.

## Contributors

N.J., A.S., F.H., and D.M.S. conceived and designed the study. A.S., S.L., D.U., and M.A.L. acquired the data and performed the formal analysis. N.J., A.S., H.S., and D.M.S. contributed to the result interpretation and critical discussion. N.J. and D.M.S. drafted the original manuscript and prepared the figures and tables. All authors contributed to the manuscript revision, reviewed the final version, and approved it for submission.

## Data sharing statement

All AIR-seq datasets used in this study are publicly available, and no individual-level data were collected specifically for this investigation. The source and access information for each dataset are detailed in Supplementary Table S1. This study was not subject to an institutional protocol and did not require participant consent. Processed donor-level feature matrices will be made available upon submission.

## Declaration of interests

A.S. and F.H. are employees and shareholders of MyImmune Corporation. D.M.S. is a shareholder in MyImmune Corporation. All other authors declare no competing interests.

## Acknowledgements

We thank all members of the Systems Immunology Lab at Osaka University for their helpful discussions and support. This work was supported by the Japan Society for the Promotion of Science (JSPS) KAKENHI Grant [JA23H034980], the Japan Agency for Medical Research and Development (AMED) Grant [JP25am0101001], and the Kishimoto Foundation Fellowship (N.J.).

## Notes

### Summary of Updates

We have corrected minor typographical and grammatical errors.

