## Supplementary for "Accurate and scalable multi-disease classification from adaptive immune repertoires"

#### Supplementary data

**Supplementary Figure S1. Confusion matrices from the remaining five cross-validation folds.** Donor-level classification results for the additional four folds corresponding to the 5-fold evaluation shown in Fig. 3c.

**Supplementary Figure S2. Donor-level classification performance on the Mal-ID cohort across different input types.** Confusion matrices are shown for 5-fold cross-validation using data from Mal-ID study. (**a**) BCR input only (n = 550). (**b**) TCR input only (n = 499). (**c**) Combined TCR and BCR input (n = 499).

**Supplementary Figure S3. t-SNE projection of encoded TCR paratope features split by disease class.** Each panel shows the same t-SNE embedding from Fig. 4a, with individual panels highlighting one of the 16 disease classes from the 1,421-donor dataset.

**Supplementary Figure S4. Heatmap of V gene usage across samples reveals study-specific clustering.** The heatmap displays the presence or absence of V genes across donor samples, highlighting that V gene usage patterns cluster strongly by study or disease cohort. Due to the large number of samples, a representative zoomed-in region is shown, illustrating that samples from the AIH dataset cluster tightly together.

**Supplementary Table S1. Immune repertoire dataset used in this study.**
The table lists all 1,421 donors across 16 disease classes, including class IDs, sample counts, and source cohort information for each group.

**Supplementary Table S2. Summary of classification performance using different input features and cohorts.**
The table summarizes the performance of our machine learning model in disease classification using AIR data from the full dataset (16 disease classes) and the MAL-ID subset (6 disease classes). The MyImmune encoder was used for TCR, while IgBERT was used for BCR data.

**Supplementary Table S3. Open-source protein language models used in our benchmarking.** The table summarizes the publicly available encoder models evaluated in this study, including their input type, model name, and a brief description of each model.

**Supplementary Table S4. Interactive t-SNE plot links for TCR datasets.**
This table provides direct links to interactive t-SNE visualizations of donor-level TCR paratope embeddings from the full 16-class dataset (1,426 donors) and the MAL-ID subset (6 classes, 499 donors).

| Disease/Health status | Count | Class | Study | Reference |
|---|---|---|---|---|
| Coronavirus disease 2019 (COVID-19) | 58 | 0 | Zaslavsky et al., 2025 (MAL-ID) | 6 |
| Healthy donors (Total = 379 donors) | 154 | 1 | Zaslavsky et al., 2025 (MAL-ID) | 6 |
| | 104 | 1 | Aterido et al., 2024 | 25 |
| | 60 | 1 | Schultheiß et al., 2021 | 23 |
| | 36 | 1 | Zuckerbrot-Schuldenfrei et al., 2024 | 17 |
| | 25 | 1 | Komech et al., 2018 | 18 |
| Human Immunodeficiency virus (HIV) infection | 95 | 2 | Zaslavsky et al., 2025 (MAL-ID) | 6 |
| Influenza vaccination recipients | 37 | 3 | Zaslavsky et al., 2025 (MAL-ID) | 6 |
| Systemic lupus erythematosus (SLE) | 63 | 4 | Zaslavsky et al., 2025 (MAL-ID) | 6 |
| Type 1 diabetes (T1D) | 92 | 5 | Zaslavsky et al., 2025 (MAL-ID) | 6 |
| Pancreatic ductal adenocarcinoma (PDAC) | 163 | 6 | Pothuri et al., 2024 | 14 |
| Non-small cell lung cancer (NSCLC) | 80 | 7 | Formenti et al., 2018 and Wu, 2021 | 15,16 |
| Ovarian cancer (OC) | 49 | 8 | Zuckerbrot-Schuldenfrei et al., 2024 | 17 |
| Ankylosing spondylitis (AS) | 44 | 9 | Komech et al., 2018 | 18 |
| Multiple sclerosis (MS) | 14 | 10 | Shugay et al., 2015 | 19 |
| Head and neck squamous cell carcinoma (HNSCC) | 56 | 11 | Luoma et al., 2022 | 20 |
| Melanoma | 38 | 12 | Robert et al., 2014 and Yusko et al., 2019 | 21,22 |
| Autoimmune hepatitis (AIH) | 59 | 13 | Schultheiß et al., 2021 | 23 |
| Nasopharyngeal Carcinoma (NPC) | 31 | 14 | Zhang et al., 2022 | 24 |
| Rheumatoid arthritis (RA) | 163 | 15 | Aterido et al., 2024 | 25 |

| | Full dataset | MAL-ID dataset | | |
|---|---|---|---|---|
| **Input type** | TCR | TCR | BCR | BCR+TCR |
| **n Class** | 16 | 6 | 6 | 6 |
| **n Donor** | 1421 | 499 | 550 | 499 |
| **Accuracy** | 0.8177 ± 0.0090 | 0.8819 ± 0.0562 | 0.8400 ± 0.0306 | 0.9039 ± 0.0327 |
| **AUROC** | 0.9885 ± 0.0012 | 0.9781 ± 0.0111 | 0.9718 ± 0.0073 | 0.9824 ± 0.01 |
| **F1 score** | 0.7929 ± 0.0177 | 0.8576 ± 0.0644 | 0.8386 ± 0.0386 | 0.8926 ± 0.0399 |

Supp. Figure S1

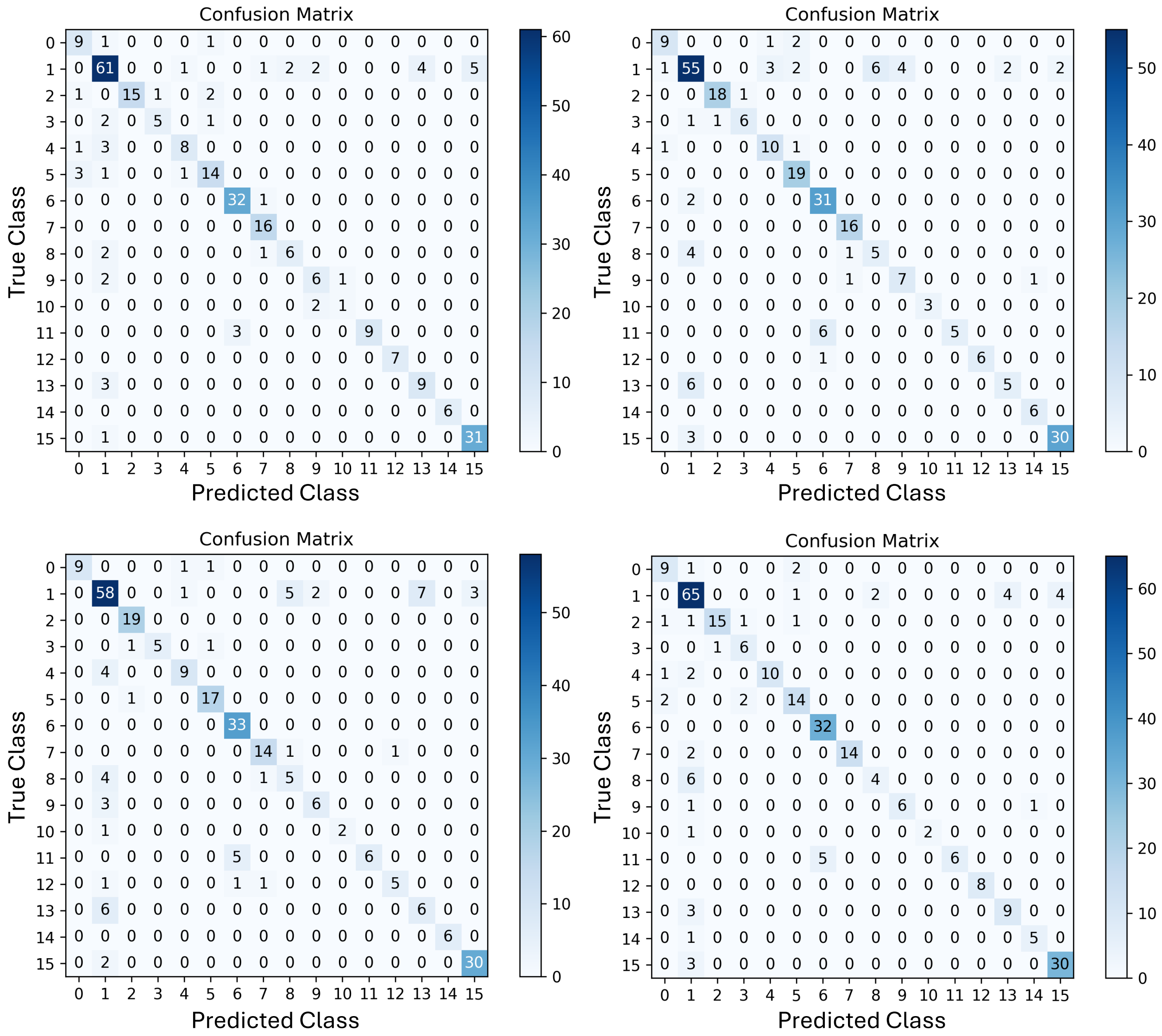

Supp. Figure S2

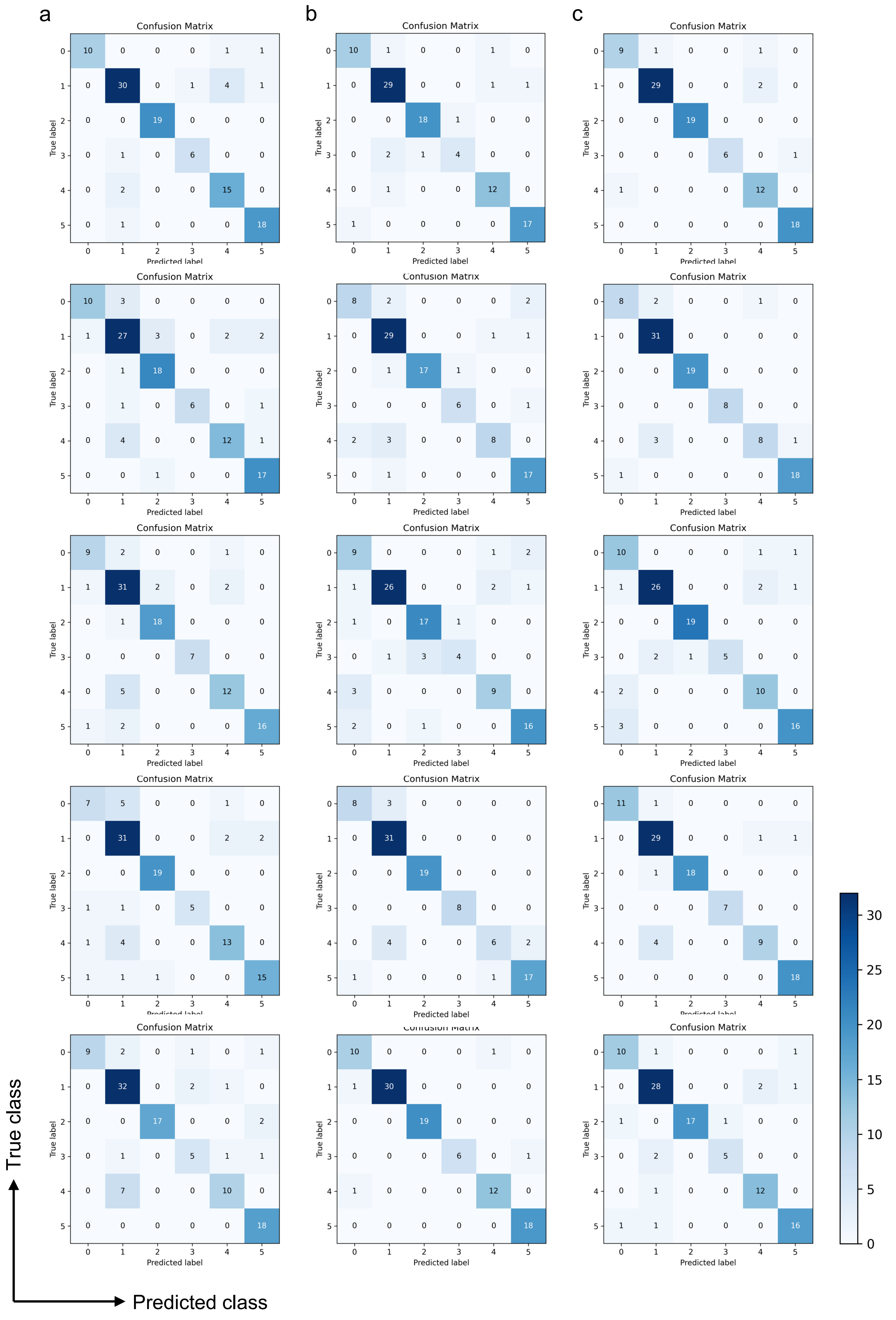

### Supp. Figure S4

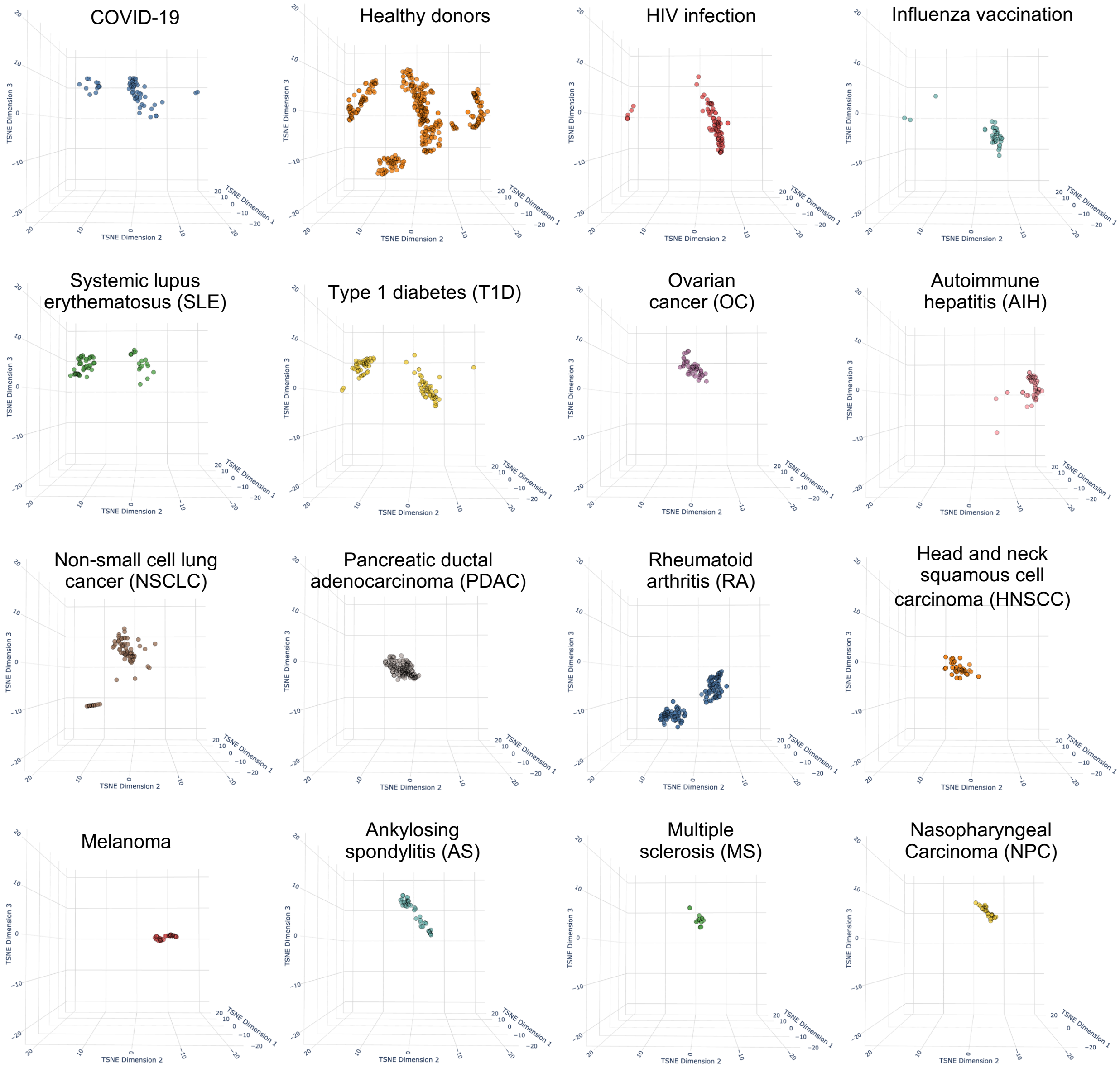

Supp. Figure S4

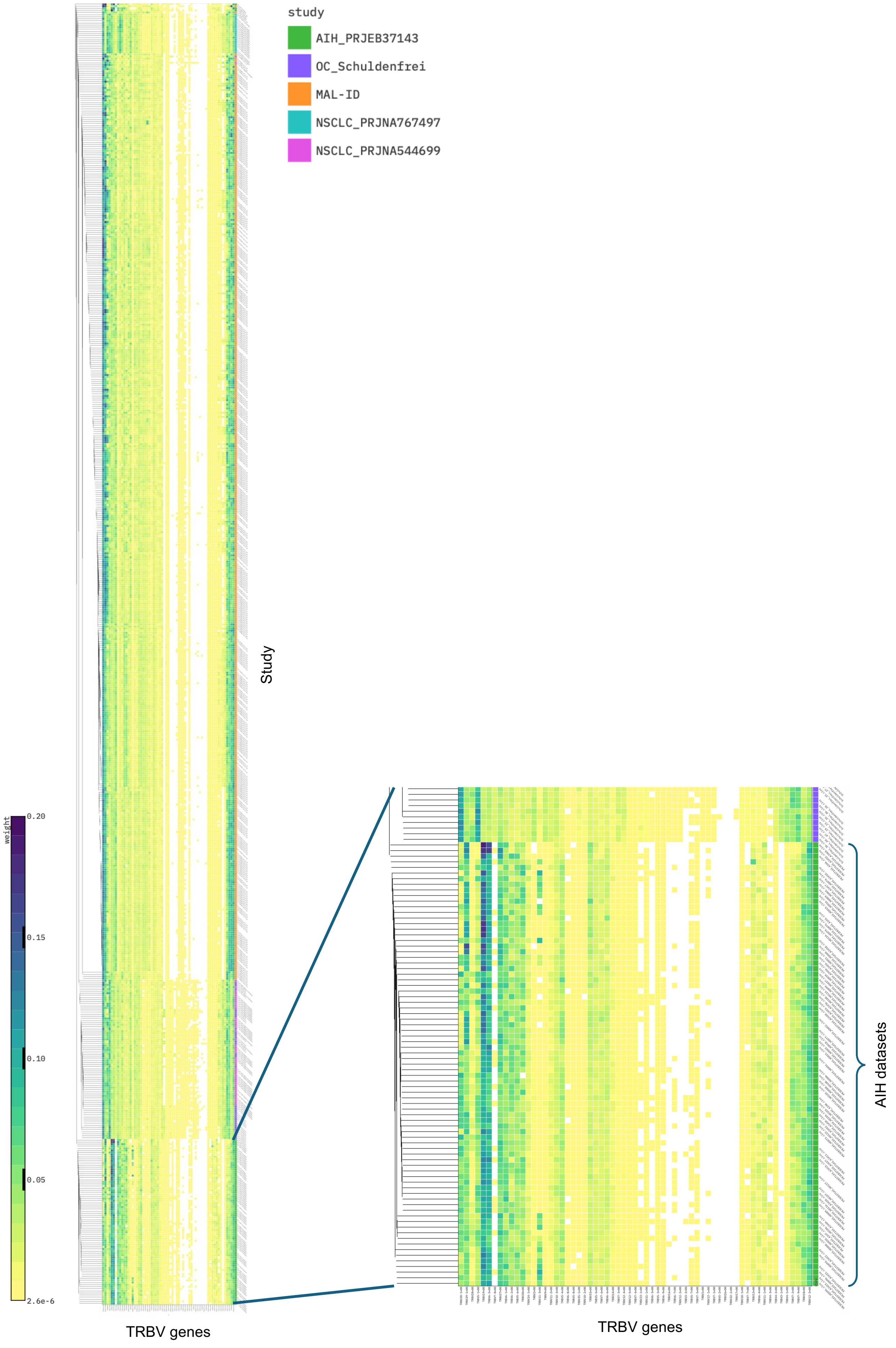

| Input type | Protein language encoders | Model | Description |
|---|---|---|---|
| Both | Evolutionary Scale Modeling version 2 (ESM2) | 8 millions, 150 million and 650 million parameters (8M, 150M, and 650M) | Evolutionary Scale Modeling version 2 (ESM2) is a family of Transformer models (ranging from 8 million to 650 million parameters) trained on UniRef50. It learns broad protein “grammar” and evolutionary covariation patterns, providing rich embeddings transferable to antibodies and other proteins. |
| Both | ProtTrans | | ProtBert is a BERT-based protein language model from the ProtTrans family, pretrained on the BFD database of over two billion sequences using masked-language modeling to learn rich, context-aware amino acid representations. |
| BCR | AntiBERTy | | A transformer-based masked-language model pre-trained on millions of human BCR sequences. It captures antibody-specific sequence patterns and supplies context-aware embeddings across framework and CDR regions. |
| BCR | AbLang | | AbLang is a BERT-style model trained solely on curated immunoglobulin heavy- and light-chain repertoires. It learns structural and functional motifs unique to antibodies through masked-language modeling. |
| BCR | IgBERT | | IgBERT is a BERT-derived model pre-trained on large datasets of immunoglobulin sequences. It includes specialized tokenization of CDR loops, enabling fine-grained representation of antigen-binding regions. |
| BCR | AntiBERTa2 | | AntiBERTa2 is an updated iteration of the AntiBERTa family with architectural enhancements and expanded training data to better capture the diversity of BCR repertoires. |
| TCR | TCRBert | | TCRBert is a BERT-based model pre-trained on large TCR-β repertoires to capture V(D)J recombination patterns and CDR3 loop diversity. |
| TCR | SCEPTR | Large model, Beta chain model, and Default model | SCEPTR is a family of Transformer models designed for TCRs, including variants for large models, β-chain-specific modeling, and default architectures. It incorporates peptide-MHC binding information into its training. |
| TCR | UniTCR | | UniTCR is a unified embedding framework that combines raw TCR sequence input with encoded V/D/J gene usage, yielding representations optimized for antigen-specificity prediction. |

| **TCR dataset** | **tSNE plot** |
| --- | --- |
| 16 classes, 1,426 donors | https://reports.myimmune.ai/tsne_tcr_16_classes/index.html |
| 6 Classes (MAL-ID cohort), 499 donors | https://reports.myimmune.ai/tsne_tcr_6_classes/index.html |
